# Habitat context alters the pace of climate-driven community warming across terrestrial and freshwater ecosystems

**DOI:** 10.64898/2026.05.06.723129

**Authors:** Emilie E. Ellis, Jussi Mäkinen, Andréa Davrinche, Irene Conenna, Laura H. Antão, Maria Hällfors, Andrea Santangeli, Benjamin Weigel, Janne Heliölä, Ida-Maria Huikkonen, Mikko Kuussaari, Aleksi Lehikoinen, Reima Leinonen, Maija Salemaa, Anna Suuronen, Tiina Tonteri, Kristiina Vuorio, Anna-Liisa Laine, Marjo Saastamoinen, Jarno Vanhatalo, Tomas Roslin

## Abstract

As global temperatures rise, ecological communities are increasingly dominated by warm-affiliated species, a process known as community warming or thermophilisation. Yet, why different taxa exhibit different rates of community warming remains unclear. Habitat composition and structure are likely drivers of this variation, as the ecological consequences of warming are filtered by local environmental conditions. Using over 40 years of monitoring data spanning terrestrial (birds, insects, plants) and freshwater (phytoplankton) communities, we show that habitat structure determines how strongly communities track warming. Forest cover systematically slows thermophilisation by reducing communities’ sensitivity to temperature change, whereas habitat heterogeneity has weak and variable effects that differ among ecosystems. Together, these results demonstrate that uneven thermophilisation arises from habitat-mediated differences in how communities respond to a shared climatic signal. Incorporating these effects is essential for improving predictions of biodiversity change under ongoing climate warming.

## INTRODUCTION

Climate change is reshaping ecological communities worldwide as species adjust their distributions and abundances in response to shifting thermal conditions (Sunde et al., 2023; Antão et al., 2020; Montràs-Janer et al., 2024). These dynamics often lead to community thermophilisation, i.e., a gradual increase in the thermal affinity of species within local assemblages, driven by warm-affiliated species colonising previously unsuitable areas or increasing in relative abundance, while cold-affiliated species retreat from or decrease within their historical ranges (Oliver et al., 2017; McLean et al., 2021; Fourcade et al., 2021; Ellis et al., 2025a). This process is commonly quantified using the Community Temperature Index (CTI) (Marjakangas et al., 2022; Khaliq et al., 2024; Devictor et al., 2008), which measures the average thermal affinity of the species in a community. A species’ thermal affinity is commonly quantified as the Species Temperature Index (the mean temperature of a species’ distribution (Schweiger et al., 2014)). The CTI is either weighted by their relative abundances to examine structural change (e.g. Åke Lindström et al. (2013)) or unweighted as a measure of turnover (e.g. Fourcade et al. (2021)). CTI has become a powerful, integrative measure of how biodiversity responds to climate warming through species turnover, colonisations, extinctions, and abundance shifts (Ellis et al., 2025a; Mäkinen et al., 2025; Devictor et al., 2008; Khaliq et al., 2024; Devictor et al., 2012).

Recent research has revealed extensive thermophilisation across terrestrial (e.g. Khaliq et al. (2024); Zellweger et al. (2020)), freshwater (e.g. Comte et al. (2021); Aalto et al. (2016); Tunney et al. (2014)), and marine systems (e.g. McLean et al. (2021); Burrows et al. (2019); Cheung et al. (2013)), including endotherms (e.g. Munter (2025); Devictor et al. (2012)), ectotherms (e.g. Ellis et al. (2025a); Devictor et al. (2012)), and plants (e.g. Govaert et al. (2021); Gottfried et al. (2012); Zellweger et al. (2020)). Yet, to date, most analyses of community warming have been taxon-specific (but see Khaliq et al. (2024); Devictor et al. (2012)), thus restricting the scope for direct comparisons across taxa. In our recent study in Finland (Mäkinen et al., 2025), we addressed this gap by conducting a cross-taxon assessment of thermophilisation trends for plant, bird, moth, butterfly, and freshwater phytoplankton communities over several decades. This study revealed divergent thermophilisation trajectories, with pronounced warming in bird, butterfly and moth communities but weaker or negligible shifts for plants and phytoplankton. Such taxon-specific responses raise a key question: *why* do communities exposed to the same climate forcing respond so differently?

A putative driver underpinning this variation is habitat context. Here, we define habitat context in terms of landscape composition and configuration. Habitat context is increasingly recognised as both a buffer (Comte et al., 2021; Zellweger et al., 2020; Fourcade et al., 2021) and an amplifier of thermophilisation (Munter, 2025; Fourcade et al., 2021). Habitat structure mediates the microclimatic conditions that organisms experience and affects dispersal and colonisation dynamics (Ash et al., 2017; Parmesan, 2006), thus shaping the persistence of cold- and warm-affiliated species (Comte et al., 2021; Fourcade et al., 2021). For example, closed-canopy forests can buffer warming effects by providing thermal refugia (Frenne et al., 2013; Zellweger et al., 2020), whereas open or degraded landscapes may accelerate community turnover toward warm-adapted species (Oliver et al., 2017). Furthermore, as these modulating effects can emerge through multiple, potentially contrasting mechanisms and species have different traits, it remains unclear whether different types of habitat composition and structure exert consistent or scale-dependent effects across taxa and ecosystems. Despite growing evidence of the role of habitat in mediating climatic effects (Frenne et al., 2019; Zhou et al., 2025; Greiser et al., 2020; Finocchiaro et al., 2024), its influence is rarely integrated in studies across taxa and/or large environmental gradients.

To determine the relative contributions of variation in local climatic exposure versus habitat-mediated differences to overall patterns of thermophilisation, we must move beyond focusing on single taxonomic groups, specific habitat types, or isolated mechanisms towards a cross-taxon and multi-habitat perspective that can assume effects through a range of mechanisms. Finland provides an exceptional setting to address these knowledge gaps. Its strong climatic gradients (Aalto et al., 2016), extensive forest cover (Korhonen et al., 2024) of varying management intensity, and detailed long-term biodiversity monitoring enable a throughout assessment of how habitat context shapes thermophilisation. Coordinated monitoring programs for birds, butterflies, moths, plants, and freshwater plankton offer decades of standardised time-series spanning broad latitudinal and environmental ranges, allowing us to thoroughly quantify temporal changes in CTI and the environmental drivers influencing them.

Here, we extended the cross-taxon analysis of (Mäkinen et al., 2025) to test how habitat composition and configuration influence the rate of thermophilisation across terrestrial and freshwater communities. Using spatial scales relevant to local communities for each taxon (Fig. S1), we quantified how changes in temperature, forest cover area, and habitat entropy (a measure of landscape heterogeneity) jointly shape temporal CTI trends. Specifically, we modelled the rate of CTI change as a function of local temperature trends and habitat characteristics, including their interactions (temperature × forest cover, temperature × entropy), to reveal how habitat composition and configuration modify the resulting community warming through space and time.

Our overarching goal was to test the buffering capacity of habitat, specifically the extent to which habitat composition and configuration modulate the strength of climate-driven thermophilisation. We address three linked questions: (1) How strongly does local temperature drive community thermophilisation across space and time? (2) Does habitat composition and configuration modify the relationship between temperature and thermophilisation, and if so, how? (3) Do taxa differ in how habitat context mediates their responses to warming, and what explains this variation? To answer these questions we analysed the spatiotemporal variation in CTI using statistical models that estimate both the overall effects of temperature and habitat covariates on CTI across space and time, and the proportion of temporal variation in CTI explained by each covariate at each monitoring site. This enables us to quantify both the strength of climate-driven thermophilisation and the relative importance of temperature versus habitat context in shaping temporal community change.

*A priori*, we set four hypotheses (Fig. 1): (H1) Ambient temperature change is the primary driver of community thermophilisation. We therefore expect a significant positive association between temperature and CTI and a high importance of local temperature trends on CTI change. (H2) Greater forest cover will buffer thermophilisation, as closed-canopy habitats reduce exposure to extreme temperatures and provide microclimatic refugia. We therefore expect a negative interaction between temperature and forest cover, resulting in weaker CTI increases in landscapes with greater forest cover. (H3) Habitat entropy, reflecting the structural and compositional diversity in the landscape surrounding each site, will have non-directional or context-dependent effects, because heterogeneous landscapes can both generate microclimatic refugia that slow thermophilisation (e.g. through more forest cover), or facilitate colonisation by warm-adapted species (e.g. more farmland or urban areas). As a result, we anticipate a mixed or taxon-specific interaction effect of temperature and entropy on CTI. (H4) Taxa differ in their responses to climate warming because they use and experience habitat differently. Taxa that are more strongly influenced by local habitat conditions and microclimates are expected to show greater modulation of thermophilisation by habitat composition and configuration, reflected in stronger interactions between temperature and habitat variables (forest cover and entropy). In contrast, taxa less dependent on local habitat conditions are expected to show more direct responses to temperature, with weaker habitat-mediated effects.

**Figure 1.**
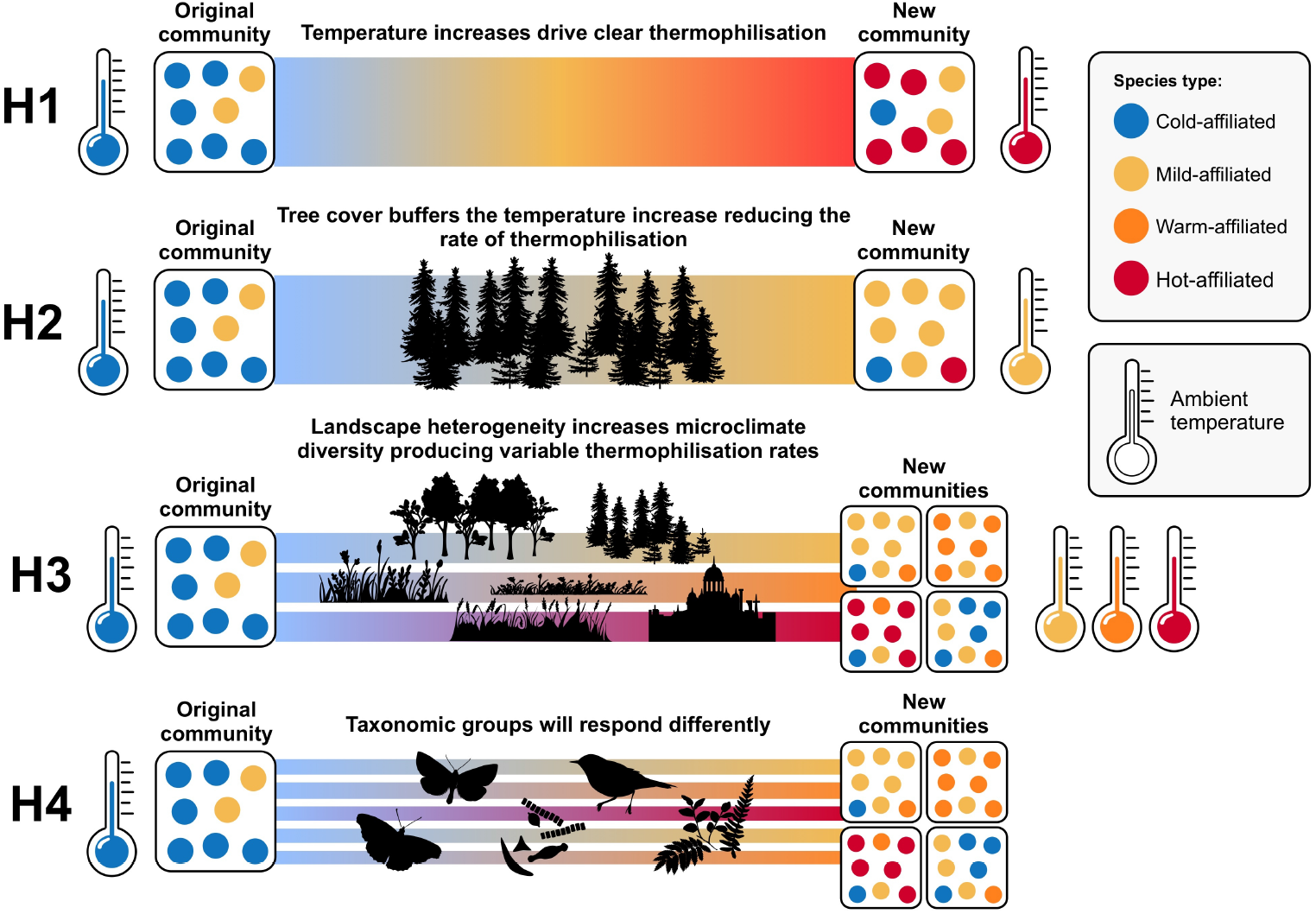
Conceptual framework illustrating four hypotheses (H1–H4) linking warming, habitat context, and community thermophilisation across terrestrial (birds, butterflies, moths and plants) and freshwater (phytoplankton) communities. Panels show shifts from an original community (left) to a new community (right) under increasing temperature. Coloured circles represent species with different thermal affinities (blue = cold-, yellow = mild-, orange = warm-, red = hot-affiliated). Thermometers and background gradients indicate increasing ambient temperature. H1: Rising temperature drives thermophilisation, with communities shifting in relative dominance/abundance from cold-to hot-affiliated species. H2: Forest cover buffers warming by moderating temperature change, thus slowing down thermophilisation. H3: Landscape heterogeneity generates diverse microclimates, producing variable thermophilisation rates within and among communities. H4: Taxonomic groups respond differently due to differences in dispersal ability, habitat dependence, and life-history traits.

## RESULTS

### Temporal patterns of environmental variables and CTI

We assessed change in temperature (Δ*°C/year*), forest cover (Δ*coverage/year*) and entropy (Δ*unit/year*) over time surrounding each monitoring site (Fig. S1). Overall, temperature changed more rapidly than habitat structure and composition (Fig. 2A). Temperature increases at bird monitoring sites were moderate and variable (mean °*C/year* = 0.009; range = −0.027 to 0.045), whereas plant monitoring sites showed stronger and more consistent increases (mean °*C/year* = 0.13; range = 0.067 to 0.196). This difference likely reflects variation in spatial coverage (Fig. 2B–F) and temporal resolution among taxonomic groups. In particular, the higher rates observed for plants are likely driven by their lower sampling frequency, with surveys conducted at decadal intervals rather than annually (see Methods). In contrast, temporal changes in forest cover and habitat entropy were minimal across sites and taxa, with slopes centred around zero (forest cover: range Δ*coverage/year* = 0 to 0.002; habitat entropy: range Δ*unit/year* = −0.002 to 0.0004).

**Figure 2.**
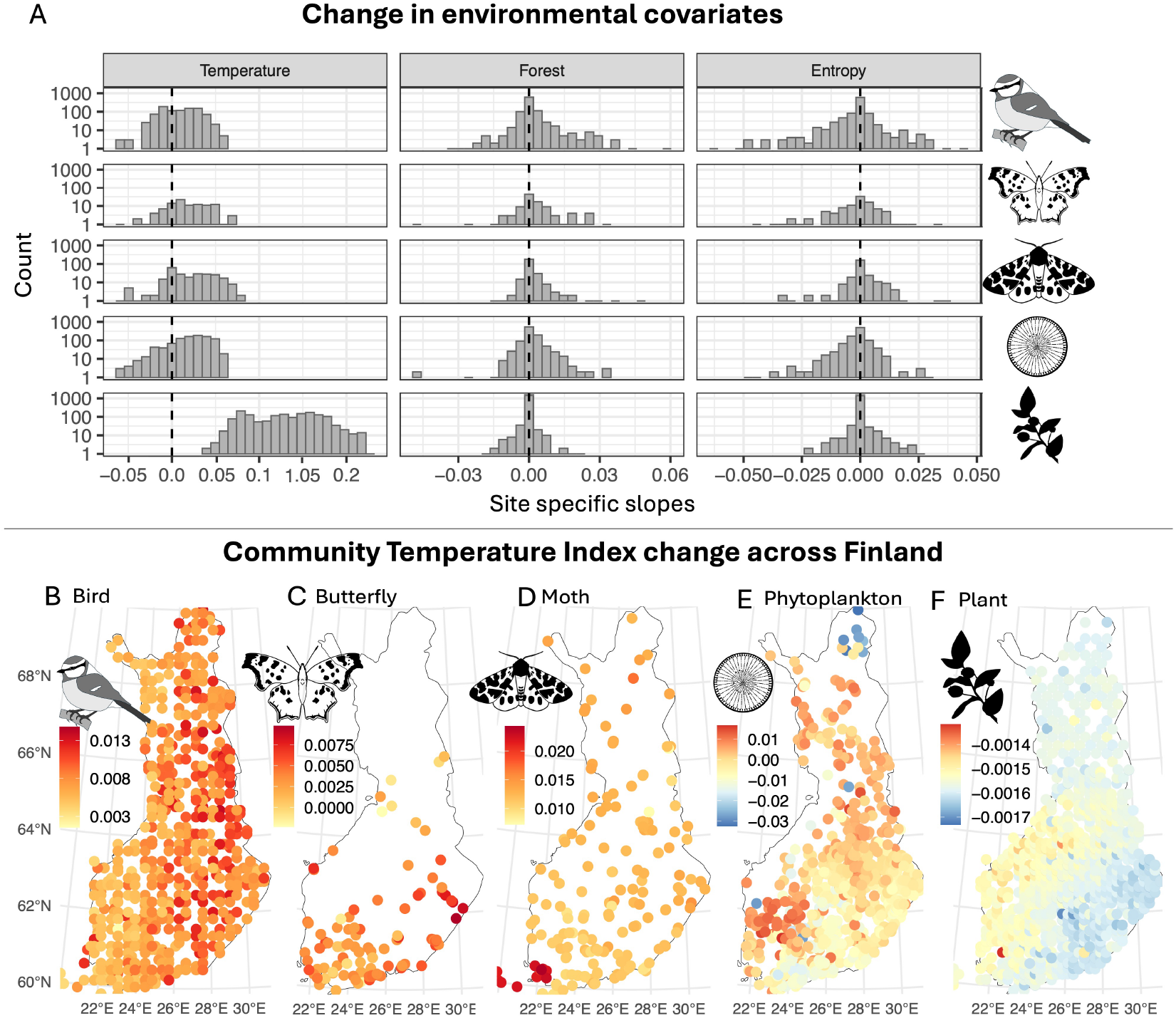
Site and taxa specific trends in environmental covariates. (A) shows the distribution of site-specific temporal slopes in temperature (Δ*C*°*/year*), forest cover (Δ*coverage/year*), and habitat entropy (Δ*unit/year*) for each taxonomic group. Histograms represent frequency distributions of site-level slopes (log_10_-scaled y-axis), with dashed vertical lines indicating 0 (no temporal change). B)-F) show taxa-specific spatial patterns through site-level temporal slopes of Community Temperature Index (CTI) change across monitoring sites in Finland; (B) birds, (C) butterflies, (D) moths, (E) phytoplankton, and (F) understory plants. CTI values were centred within each taxon prior to modelling. Slopes represent posterior mean site-level random slopes for year extracted from the hierarchical model and indicate rates of change in CTI (centred units per year). Warmer colours indicate positive slopes (thermophilisation), cooler colours indicate negative slopes. Colour scales differ among panels to reflect taxon-specific ranges in CTI change Δ*CTI/year*.

We show that site-level CTI trends differed among taxonomic groups (Fig. 2B-F). Overall, birds, moths, and butterflies showed increases in CTI over time, indicating community thermophilisation across most sites (mean slopes Δ*CTI/year*: birds = 0.0076, 95% interval [0.0043, 0.0124]; moths = 0.0044, 95% interval [0.0090, 0.0221]; butterflies = 0.0036, 95% interval [−0.0013, 0.0080]). In contrast, plant communities exhibited CTI trends that were consistently slightly negative but close to zero across sites (mean = −0.0016, 95% interval [−0.0016, −0.0015]). Phytoplankton showed highly variable site-level trajectories centred near zero (mean = −0.0022, 95% interval [−0.0153, 0.0110]).

### Environmental drivers of thermophilisation

Temperature emerged as the strongest and most consistent predictor of CTI across all taxa (Fig. 3A). Because all variables were standardised prior to analysis, coefficients represent the change in one-standard deviation of CTI associated with a one–standard deviation increase in each covariate. Temperature increase was associated with marked increases in CTI in moths (mean Δ*CTI/year* = 0.81; range = 0.79, 0.83), birds (mean Δ*CTI/year* = 0.71; range = 0.69, 0.73) and butterflies (mean Δ*CTI/year* = 0.46; range = 0.41, 0.51), but weaker CTI increases for plants (mean Δ*CTI/year* = 0.24; range = 0.21, 0.27) and phytoplankton (mean Δ*CTI/year* = 0.24; range = 0.21, 0.27).

**Figure 3.**
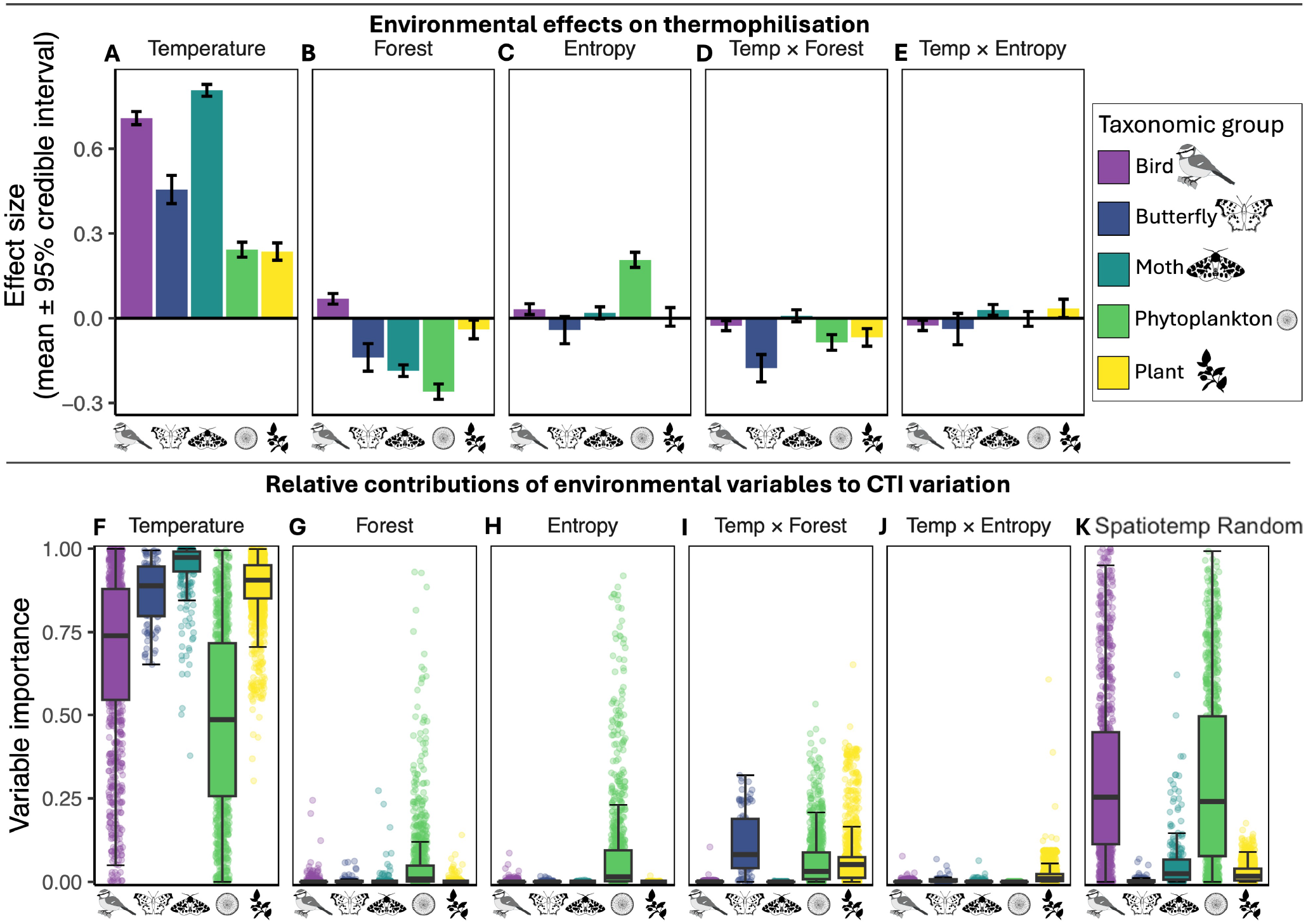
Effect size estimates and variance components from models relating temperature change and habitat context to change in community temperature index (CTI) across five taxonomic groups in Finland. All predictors were standardised prior to analysis; effect sizes therefore represent the change in CTI associated with a one standard deviation change in each covariate. A–E, estimated effects (mean ± 95% credible intervals) of (A) temperature, (B) forest cover, (C) habitat entropy, (D) temperature × forest cover interaction, and (E) temperature × entropy interaction on change in CTI. Positive values indicate increases in CTI, whereas negative values indicate decreases. F–K, variance partitioning showing the proportion of temporal variance in CTI per monitoring site attributable to (F) temperature, (G) forest cover, (H) habitat entropy, (I) the temperature × forest cover interaction, (J) the temperature × entropy interaction, and (K) the spatio-temporal random effect representing residual spatial and temporal structure not explained by the covariates. Distributions reflect the variation of variance partitioning across monitoring sites and are shown as boxplots (median, interquartile range, and range), with points representing individual site-level estimates. Colours and icons denote taxonomic groups.

Habitat structure and composition had a direct effect on CTI in all taxonomic groups (Fig. 3B). Forest cover had a negative effect on CTI in moths (Δ*CTI/year* = −0.19; range = −0.21, −0.17), butterflies (Δ*CTI/year* = −0.14; range = −0.19, −0.09), plants (Δ*CTI/year* = −0.04; range = −0.07, −0.01), and phytoplankton (Δ*CTI/year* = −0.26; range = −0.29, −0.24), suggesting slower thermophilisation in more forested sites. Birds were the only group showing a positive effect of forest cover on CTI (Δ*CTI/year* = 0.07; range = 0.05, 0.09), i.e. higher CTI was associated with higher forest cover. Effects of habitat entropy were weaker and less consistent across taxa (Fig. 3C). Entropy had a positive effect on CTI in phytoplankton (Δ*CTI/year* = 0.21, range = 0.18; 0.27) and birds (Δ*CTI/year* = 0.03, range = 0.01; 0.05), whereas the effects were weak or uncertain in moths (Δ*CTI/year* = 0.02; range = 0.003, 0.04), plants (Δ*CTI/year* = 0.005; range = −0.03, 0.04), and butterflies (Δ*CTI/year* = −0.04; range = −0.09, 0.006).

Interactions between temperature and habitat variables further indicated that habitat context mediated CTI-responses to warming (Fig. 3D,E; Fig. 4). The interaction between temperature and forest cover was negative in birds (ΔCTI = −0.03; range = −0.04 to −0.01), butterflies (ΔCTI = −0.18; range = −0.23. −0.13), plants (ΔCTI = −0.07; range = 0.10, −0.04), and phytoplankton ((ΔCTI = −0.03; range = −0.06, −0.0001), indicating that forest cover dampened temperature-driven increases in CTI (i.e. thermophilisation sensitivity, Fig. 4A). For moths the interaction between temperature and forest cover overlapped zero.

**Figure 4.**
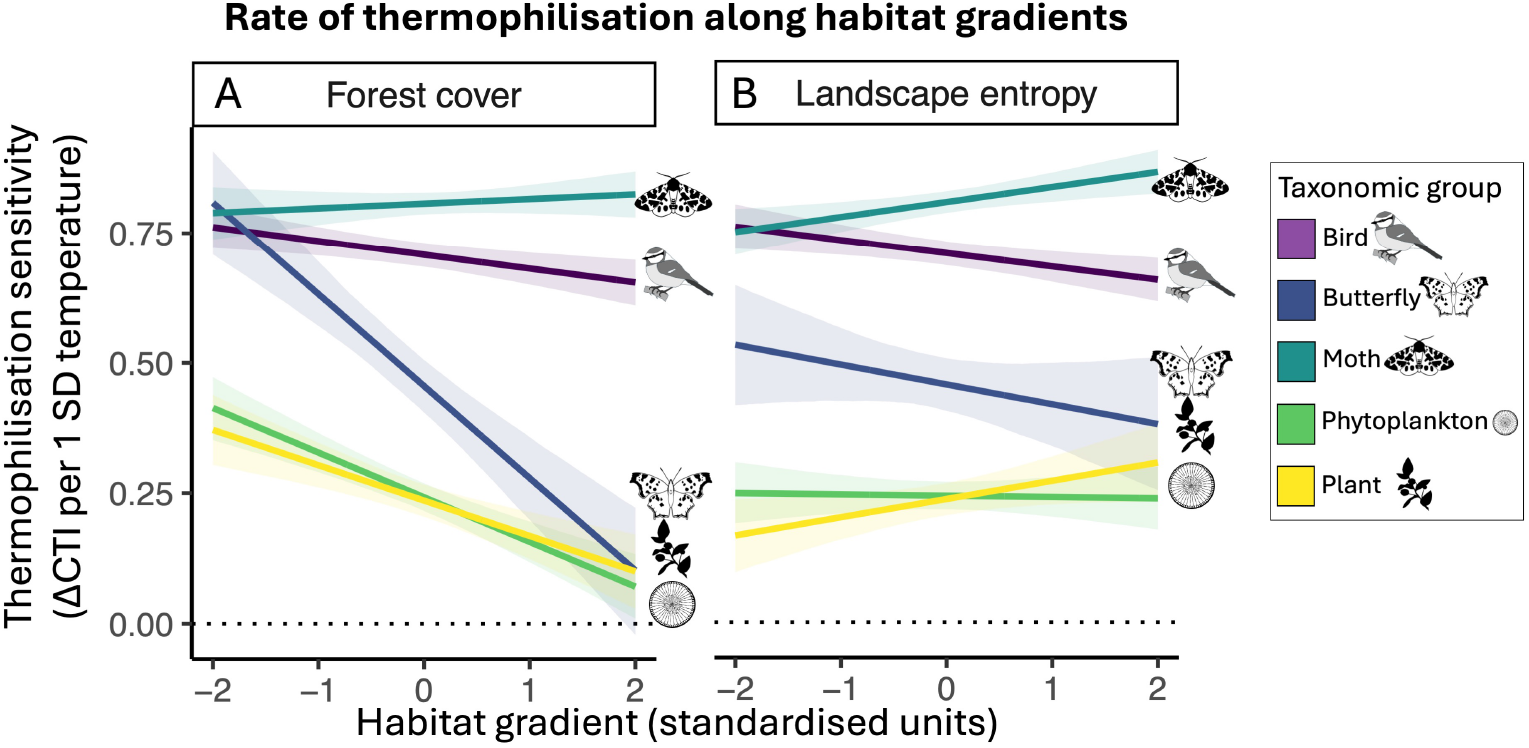
Predicted temperature sensitivity of the Community Temperature Index (CTI) across gradients of forest cover and landscape entropy for birds, butterflies, moths, phytoplankton, and plants. Temperature sensitivity was calculated from fitted interaction models as the expected change in CTI associated with a one-standard-deviation increase in temperature, evaluated across standardised habitat gradients from −2 to +2 SDs. Predictions, shown as coloured lines, were generated from the temperature main effect and the interaction between temperature and (A) forest cover or (B) entropy while holding the other habitat context variable constant. Shaded ribbons show approximate 95% uncertainty intervals derived from posterior standard deviations of the corresponding coefficients. The horizontal dotted line at zero indicates no temperature sensitivity of CTI.

The interaction between temperature and habitat entropy were generally weak and taxon-specific (Fig. 3 E; Fig. 4B), entropy was associated with lower thermophilisation sensitivity in birds but higher thermophilisation sensitivity in moths and plants. Though these effects were close to zero and there no clear effect on thermophilisation sensitivity in butterflies or phytoplankton.

### Variance partitioning of environmental drivers

Variance partitioning shows the extent that the covariates and spatio-temporal random effect contribute to the temporal variation of CTI at each monitoring site. The variance partitioning results supported the patterns reported above (Fig. 3): temperature was the dominant environmental driver of temporal variation of CTI across all taxa (Fig. 3F). Temperature accounted for the largest share of CTI variation in terrestrial communities, representing 94% of variable importance in moths, 89% in plants, 87% in butterflies, and 68% in birds. In contrast, its contribution was substantially lower in phytoplankton (49%), indicating weaker coupling between local temperature trends and CTI dynamics in freshwater communities.

Habitat variables explained comparatively little temporal variation of CTI but showed taxon-specific patterns. Forest cover (Fig. 3G) contributed minimally to terrestrial taxa (*<*1% in birds, butterflies, moths, and plants), but slightly more to phytoplankton (5%). Habitat entropy (Fig. 3H) contributed little to temporal variation of CTI in terrestrial groups (*≤*1%) but more in phytoplankton (9%). The interaction between temperature and forest cover contributed 12% of temporal variation of CTI for butterflies, 7% for plants, and 6% for phytoplankton (Fig. 3I), but was negligible for birds (0.1%) and moths (*<*0.1%). The interaction between temperature and entropy had generally low importance across all taxa (*≤*1%). Compared to the relatively large effect estimates of interactions, forest cover and entropy have a small role as the driver of temporal variation CTI. Low importance of forest and entropy on temporal variation of CTI can be explained by the relatively small temporal change in them (Fig. 2A). For birds and phytoplankton, the spatio-temporal random effect (Fig. 3K) explained a substantial share of CTI variation (31.7% and 30.9%, respectively), indicating considerable inter-annual variability not captured by the measured environmental predictors. The importance of covariates and spatio-temporal random effect in variance partitioning varied strongly between monitoring sites, highlighted by the site-level values in Fig. 3F-K. This again reflects the importance of site-level context for the drivers of CTI dynamics.

## DISCUSSION

Our results show that temperature exerts a consistent positive effect on CTI across systems. However, habitat context also plays a role in shaping how closely ecological communities track ambient temperatures. Across the five major taxonomic groups spanning terrestrial and freshwater systems, habitat variables not only had direct effects on CTI but also modified the strength of the temperature–CTI relationship. Below, we discuss these findings in relation to our original hypotheses (Fig. 1).

### Ambient temperature change is the primary driver of community thermophilisation

Consistent with our first *a priori* hypothesis, increasing temperature was strongly associated with increasing CTI across terrestrial taxa, with directional shifts in community thermal composition being tightly coupled with local warming. Since temperature emerged as the most rapidly changing covariate tested, this result may seem unsurprising at first. Nonetheless, the relationship between temperature and thermophilisation breaks down when moving from terrestrial to freshwater systems. Despite comparable climatic warming across lake monitoring locations (Fig. 2A), for phytoplankton, temperature had relatively low importance for explaining CTI variation and showed weak effects on expected thermophilisation (Fig. 3A,F). This divergence highlights an important constraint to cross-ecosystem generalisations. A similar limit to cross-ecosystem generality has been reported in recent syntheses (Khaliq et al., 2024).

Together, these results emphasise that similar macroclimatic warming will not necessarily translate into equivalent thermal community reassembly across ecosystems. In aquatic systems, air temperature alone may not capture the realised thermal and resource environment structuring freshwater communities (Mouton et al., 2022; Ovaskainen et al., 2019). For phytoplankton communities in particular, climate effects are often determined through indirect pathways, since warming alters the timing and strength of stratification patterns, nutrient recycling, trophic dynamics, and light availability (Adrian et al., 2009; Sommer et al., 2012; Winder and Schindler, 2004). These indirect processes can decouple community reassembly from trends in air temperature even when surface waters warm (Litchman et al., 2010; Soranno et al., 2014; Weigel et al., 2023). Furthermore, a substantial fraction of variation in phytoplankton CTI was attributed to the spatio-temporal random effect (Fig. 3K), indicating pronounced inter-annual and among-site variability not captured by change in mean temperature or change in the habitat context variables tested here. These results align with previous phytoplankton analyses (Weigel et al., 2023; Ovaskainen et al., 2019) showing that even when detailed limnological predictors are included, temporal random effects remain large, possibly because communities are predominantly structured by resource competition (Tilman, 1986) and short-term environmental variability (Winder and Schindler, 2004). The weak effect of temperature on phytoplankton communities therefore does not necessarily imply climatic insensitivity, but rather that the relevant climatic pathway for phytoplankton differs from the macroclimatic predictors that explain terrestrial thermophilisation.

Like phytoplankton, temperature had a relatively small effect size on the CTI of plant communities (Fig. 3A), indicating weak rates of thermophilisation. However, temperature consistently showed high variable importance (Fig. 3F), demonstrating that even weak temperature responses can generate a pervasive directional signal. The small magnitude of these shifts likely reflects strong ecological constraints. Because understory plants are sessile, their distributions and abundances are tightly constrained by local edaphic conditions, including soil properties, nutrient availability, and stand structure, which can limit community reassembly (Chelli et al., 2024; Kaarlejärvi et al., 2021; Majasalmi and Rautiainen, 2020). Consistent with this, Villén-Peréz et al. (2020) showed that plant responses to temperature across Finland are strongly mediated by soil and stand properties. The relatively low importance of measured environmental covariates, together with substantial spatiotemporal random effects (Fig. 3K), further suggests that unmeasured local conditions constrain the magnitude and spatial expression of change. Together, these results indicate that plant communities track warming in a consistent but constrained manner, with temperature setting the direction of change and local environmental filtering determining its magnitude.

### Forest cover alters thermophilisation rates

Consistent with our second hypothesis, forest cover influenced how closely communities track ambient temperatures. Greater forest cover was associated with lower CTI for all taxa except birds, indicating that communities embedded in more extensively forested landscapes tend to have lower thermal affinities than those in less forested landscapes under similar climatic conditions. However, compared to temperature, forest cover contributed relatively little to temporal thermophilisation, likely because forest cover changed only weakly over the study period.

The strong negative interaction between temperature and forest cover observed for butterflies, plants, phytoplankton, and birds indicates that increasing forest cover dampens the positive effect of temperature on CTI (i.e. reduces thermophilisation sensitivity, Fig. 4A). This pattern is consistent with microclimatic buffering by closed-canopy habitats: forests reduce thermal extremes (Finocchiaro et al., 2024), maintain cooler and more stable understory conditions (Greiser et al., 2020; Zhou et al., 2025; Frenne et al., 2019), and can decouple local microclimates from macroclimatic trends (Zhou et al., 2025; Frenne et al., 2013). By moderating the rate at which local thermal environments track regional warming, forested landscapes likely slow the replacement of cold-affinity species by warm-affinity species, thereby reducing rates of community thermophilisation (Fourcade et al., 2021; Lenoir et al., 2017; Zellweger et al., 2020). This buffering capacity provides a mechanistic explanation for the widely observed lag between climatic warming and community thermophilisation (Mäkinen et al., 2025; Devictor et al., 2008), indicating that forested landscapes can function as microrefugia that prolong the persistence of cold-affinity assemblages under ongoing climate change (Zhou et al., 2025; Finocchiaro et al., 2024).

Birds diverged from this general pattern in the main effect of forest cover, exhibiting higher CTI in more forested landscapes. This contrast likely reflects differences in how relatively mobile taxa respond to habitat structure. Increasing forest cover may enhance habitat availability and connectivity for warm-affinity forest generalist birds expanding northward (Lehikoinen and Virkkala, 2016), facilitating their colonisation and persistence within managed forested landscapes. This can elevate CTI through compositional changes driven by immigration and population growth of warm-adapted species (Yeakel, 2024; Munter, 2025). At the same time, the negative interaction between temperature and forest cover indicates that forests still buffer local temperature tracking, slowing the rate at which communities respond to warming. Birds therefore exhibit a combination of habitat-driven increases in CTI and microclimatic buffering of temporal responses. We note that our land-use data (CORINE) primarily captures managed and early successional forests rather than old-growth systems, which may differ in their capacity to support cold-adapted species (Santangeli et al., 2017). Together, these results suggest that thermophilisation in birds reflects both landscape-mediated species turnover and microclimatic buffering, consistent with the high importance of spatiotemporal random effects for temporal thermophilisation (Fig. 3K).

Moths represented a partial exception to the microclimatic buffering mechanism. Although forest cover had a negative effect on CTI for moths, the absence of a clear interaction between temperature and forest cover indicates that forest cover does not substantially mediate the temperature–CTI relationship in moth communities. One explanation may lie in the predominantly nocturnal activity of moths. Forest canopies strongly buffer daytime thermal extremes (Zhou et al., 2025; Finocchiaro et al., 2024), but a recent global synthesis suggests that they may also elevate nocturnal below-canopy temperatures by trapping longwave radiation (Reek et al., 2026; Geiger et al., 2009). As a result, moth communities may experience relatively warmer conditions within forests compared to diurnal taxa, weakening the capacity of forest cover to mediate temperature effects. An additional explanation may lie in resource availability and continuity. Forested landscapes can support stable and diverse moth communities by providing host plants, nectar resources, and structural complexity (Ellis and Wilkinson, 2021; Ellis et al., 2025b, 2023; Fuentes-Montemayor et al., 2022; Merckx et al., 2012). Such resource-mediated stability may maintain both cold- and warm-affinity species, thereby dampening compositional turnover even as temperatures rise. Because thermophilisation in moths is strongly linked to species pools and relative abundances (Ellis et al., 2025a), this stabilising effect may lead to a decoupling between habitat-driven and temperature-driven influences on community composition.

### Habitat entropy has weak and inconsistent effects on thermophilisation rates

Habitat entropy did not show a consistent effect on CTI across taxa, indicating that landscape heterogeneity alone does not systematically influence community thermal composition. Likewise, interactions between temperature and entropy were generally weak, suggesting that heterogeneous landscapes neither consistently amplify nor buffer thermophilisation (Fig. 4B).

A likely explanation is that entropy primarily increases habitat diversity and niche availability, promoting the coexistence of species with contrasting thermal affinities rather than directional shifts in community composition. This can elevate within-community variation without substantially altering the community-weighted thermal signal captured by CTI. In addition, the processes associated with entropy—such as fragmentation, disturbance, and resource heterogeneity—can exert opposing effects on different species, leading to weak net changes in thermal composition at the community level (Fahrig, 2003; Bogaert et al., 2005; Álvarez et al., 2024).

Phytoplankton constituted a notable exception, with higher entropy associated with increased CTI. However, because overall CTI trends in lakes were weak and slightly negative, this pattern likely reflects broader community reorganisation rather than direct thermal responses (Khaliq et al., 2024). In freshwater systems, entropy likely captures catchment-scale processes that influence phytoplankton communities, including nutrient inputs, runoff, and pollution (Dory et al., 2024; Aura et al., 2020; Winder and Sommer, 2012). This suggests that entropy operates as a proxy for watershed dynamics rather than a direct driver of thermal restructuring. Consistent with this, entropy and forest cover showed higher importance for phytoplankton than for other taxa, likely because both variables integrate multiple catchment-level processes in aquatic systems.

Overall, these results indicate at the spatial grain considered here, habitat entropy’s net effect on CTI is weak relative to the more direct and interpretable influences of temperature and forest cover.

### Ecological differences among taxa shape habitat and temperature mediated responses

Our fourth hypothesis predicted that habitat context would influence CTI in a taxon-specific manner, and we found partial support for this prediction. Across all groups, temperature acted as a consistent driver of thermophilisation, indicating a shared climatic signal. In contrast, taxa differed in the extent to which habitat buffered or facilitated temperature-driven community change, reflecting differences in how species interact with and experience their environment. For taxa closely tied to local conditions, habitat structure constrained community reassembly and dampened thermophilisation, whereas for more mobile taxa, habitat influenced colonisation dynamics and could accelerate or redirect compositional change. These results suggest that variation in thermophilisation rates among taxa arises not from fundamentally different responses to temperature, but from differences in how habitat mediates exposure to and tracking of climatic change. Community warming therefore reflects a common climatic forcing, whose expression is shaped by taxon-specific ecological interactions with habitat.

## Conclusions

Our synthesis adds a mechanistic dimension to a growing body of research seeking to understand uneven thermophilisation across ecosystems (Mäkinen et al., 2025; Khaliq et al., 2024), and clarifies how climate-driven community change unfolds within real-world landscapes. In doing so, we show that, across taxa, climate sets the direction of change, but habitat determines how it unfolds. Previous work from Finland has shown that habitat composition is a dominant determinant of *static* species occurrences and community structure (Guilbault et al., 2025, 2026). Here, we extend that framework by demonstrating that once habitat has filtered the species pool, temperature and its interactions with habitat context together govern directional thermal reassembly *through time*. Our findings reveal the complex thermal restructuring of co-occurring communities exposed to both climate change and varying habitat structure and composition, providing a pathway for integrating habitat-mediated climate responses into predictions of biodiversity change under global change.

## MATERIAL AND METHODS

To quantify the environmental drivers of community warming, we combined spatially extensive, multi-decadal monitoring data from five major taxonomic groups in Finland: birds, butterflies, moths, understory forest plants, and freshwater phytoplankton (Fig. 2). For each species, we calculated the Species Temperature Index (STI: average temperature across their European range distributions) and used these values to derive the community-level Community Temperature Index (CTI), defined as the abundance-weighted mean STI of each community at each site and each year. Plants were surveyed in 1985, 1995 and 2006 and thus the plant data corresponds more to three snapshots than temporally continuous data that we have for other species groups. At each monitoring site, we calculated the annual mean temperature and quantified surrounding habitat conditions at taxon-relevant spatial scales, including forest cover and habitat entropy, which reflects habitat diversity and evenness (Carranza et al., 2007). We examined how CTI changed through time across taxonomic groups and tested the relative contributions of ambient temperature, forest cover, and entropy, as well as their interactions, to changes in CTI.

### Species data, Species Temperature Index and Community Temperature Index

All species data and STI calculations are described in (Mäkinen et al., 2025).

In brief, we used species data from long-term Finnish monitoring programmes that cover the entire country (60°N–70°N, 19.4°E–31°E, Fig. 2). All taxonomic groups were surveyed using taxon-specific standardised protocols. Detailed descriptions of survey methods are available in the following sources: birds (Lehikoinen, 2013; Virkkala and Lehikoinen, 2014), butterflies (Heliölä et al., 2022; Kuussaari et al., 2007), moths (Leinonen et al., 2016, 2017; Huikkonen et al., 2024), plants (Reinikainen et al., 2000; Mäkipää and Heikkinen, 2003), and phytoplankton (Weigel et al., 2023).

For birds, we used records collected during 1978–2020, including 143 species sampled across 940 transects (Fig. 2A). For butterflies, records were collected during 1999–2020 and included 90 species sampled across 101 transects (Fig. 2B). For moths, records were collected during 1993–2022 and included 722 species sampled in 246 traps (Fig. 2C). Phytoplankton records include 706 species, collected between 1978-2017 in 957 study sites (Fig. 2C). For plants, 1538 quadrats of 2 m^2^ each were monitored in 1985-1986 and 1995, with a resurvey of 439 sites in 2006 (Fig. 2). The data covered 348 plant species in total. In each square, surveyors recorded the proportional coverage of vascular plants which we used as our abundance measure for weighting the STIs and produce a CTI measure. For full details on the taxon-specific data see (Mäkinen et al., 2025).

The derivation of STIs followed the general procedure outlined in (Mäkinen et al., 2025). In brief, we used each species’ European distribution to measure their thermal niche. Following the approach in Schweiger et al. (2014), we extracted range polygons from European atlas sources for birds (BirdLife’s range map catalogue BirdLife International and Handbook of the Birds of the World (2018)), butterflies (range maps derived from Schweiger et al. (2014)), and moths (range maps digitised from (Fibiger, 1990, 1993, 2009; Fibiger and Hacker, 2007; Fibiger et al., 1995, 2010; de Freina and Witt, 1987, 1990; Goater et al., 2003; Hacker et al., 2002; Hausmann et al., 2014; Hausmann and Viidalepp, 2012; Hausmann, 2004; Mironov et al., 2003; Müller et al., 2019; Ronkay and Ronkay, 1994; Ronkay et al., 2012, 2001; Skou and Sihvonen, 2015; Zilli et al., 2005)). Plant distributions were compiled from multiple continental plant atlases (Caudullo et al., 2017; Vangansbeke et al., 2021; Kalwij et al., 2014; Kurtto et al., 2018), and phytoplankton occurrences were sourced from the WISER database, which aggregates monitoring data from 26 European countries (Moe et al., 2012) to produce probabilistic range maps for these species.

All spatial data were harmonised to the 50 x 50 km Common European Chorological Grid System (CGRS). We chose to produce a baseline estimate of species’ thermal associations independent of the subsequent warming period that may be captured by the monitoring data. Therefore, each grid cell was annotated with long-term mean annual temperature for the 1961–1990 climatological reference period using the gridded climatology by Fronzek et al. (2011). Because phytoplankton lack continuous range maps, we created probabilistic range maps with species distribution models. We fitted species distribution models using European wide monitoring data of phytoplankton (starting in 1980) and explained their occurrence and predicted European ranges using mean annual temperature, annual temperature range, annual precipitation sum, and annual precipitation range (Fronzek et al., 2011). For each study species, we calculated the STI by taking the presence probability weighted mean of the mean annual temperature over all cells in CGRS.

Once we had distribution maps for each species, the STI was calculated as the mean long-term temperature across all CGRS cells weighted by the predicted occurrence probability of the species.

Community temperature indices (CTI) were then calculated at each monitoring site as the abundance-weighted mean STI of all species recorded in that assemblage:

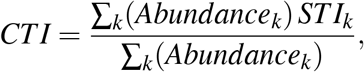

where *STI*_*k*_ is the Species Temperature Index of species *k*, and *w*_*k*_ is its abundance in the community (or proportional cover for plants).

We used the rate of the temporal change of CTI estimated in (Mäkinen et al., 2025) in Figure 2.

### Environmental data

We derived mean annual temperature data for the sampling sites based on daily records of mean temperature provided at a resolution of 10 *×* 10 km by the Finnish Meteorological Institute (Aalto et al., 2016).

Land cover data was derived from the Corine Land Cover of 2000, 2006, 2012 and 2018, which classifies land cover into 44 classes at a resolution of 25 *×* 25 m across Europe (European Environment Agency, 2000, 2006, 2012, 2018). To characterise the land cover around the monitoring sites, we followed Guilbault et al. (2026) and defined a 500 m buffer around the site for moths, butterflies, and plants and a 1 km buffer for birds (reflecting their higher mobility, Fig. S1). For phytoplankton, we used catchment area deliniations to define the area associated with each lake (Fig. S1; Röman et al. (2018)). The catchment area shapefiles were extracted from the Finnish River Basin system made by the Finnish Environment Institute (Syke; https://ckan.ymparisto.fi/dataset/valuma-aluejako). We used “level-4” of this map data to define catchment areas as it is the most accurate level of river basin division based on a nationally uniform river basin hierarchy.

To characterise land cover, buffers were drawn around each site-specific coordinate. The site coordinates were set to the centre point of a monitoring transect for birds and butterflies, to the trap location for moths, to the centre point of the plant survey site, and to the centre of the lake of phytoplankton sampling (Fig. S1). Inside each buffer, we defined the proportion of forest cover as the summed coverage of broad leaf (class 311 in Corine Land Cover data set), coniferous (class 312), and mixed forests (class 313).

To measure the diversity in land cover we used the concept of entropy, which quantifies the evenness of different land cover types (Carranza et al., 2007). Entropy was defined over all land cover classes inside the buffer:

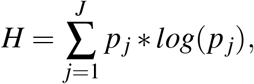

where *H* is entropy, *j* is a subindex for a land cover class, *J* is the number of different land cover classes inside the buffer and *p* is the proportion of coverage of the land cover class inside the buffer. Since species monitoring of all taxa had started before the first land cover survey (2000), we assigned land cover variables as NA for all monitoring sites prior to 2000. From 2000 onward, we assigned a linearly interpolated value of the land cover values for each year that falls between the survey years (2000, 2006, 2012, and 2018). For the years 2019-2022 we fixed the values to those observed during the 2018 survey.

### Statistical model

#### Temporal change of environmental variables

We estimated the annual rates of change in environmental drivers with linear mixed effects models. To account for variation in starting years of monitoring and in the duration of the surveys across monitoring sites, we defined the temporal responses as a site-level random effect. This allows responses to vary across sites and smooths out site-level differences (which partially originate from different sampling periods).

The linear predictor for each response variable was defined as

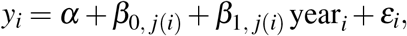

where observation *i* belongs to site *j*(*i*). Here *y*_*i*_ is the response for observation *i, α* is a global intercept, *b*_0, *j*(*i*)_ is a site-specific (random) intercept, *b*_1, *j*(*i*)_ is a site-specific (random) slope for year, and *ε*_*i*_ is an error term.

We kept the response variables within their original ranges and set the explanatory variable for year to start from zero. Outputs of the models are included in Supplementary Material (Tables S1-5).

#### Environmental drivers of CTI

We estimated the effects of environmental covariates on CTI with a linear mixed effects model,

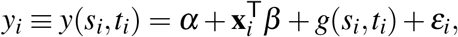

where observation *i* is taken at spatial location *s*_*i*_ and time *t*_*i*_, and **x**_*i*_ ≡ **X**(*s*_*i*_, *t*_*i*_) is the vector of covariates evaluated at (*s*_*i*_, *t*_*i*_). Here *α* is an intercept, *β* is the regression coefficient, *g*(*s*_*i*_, *t*_*i*_) is a spatio-temporal random effect, and *ε*_*i*_ is an error term.

We defined the spatio-temporal random effect *g*(*s, t*) as Gaussian processes. We applied Stochastic Partial Differential Equations (SPDE) to approximate and calculate it (Lindgren et al., 2011; Miller et al., 2019). The spatial component of random effects *g*(*s, t*) was approximated by coarsening its spatial resolution to 119 (butterflies) and to 802 (birds) tessellations. We defined the full covariance as a separable spatio-temporal structure (Cameletti et al., 2012). The covariance is a Kronecker product of spatial and temporal covariance matrices. Spatial covariance is defined with a Matérn (*α*=3/2)-type covariance function. The temporal covariance with respect to the year was defined with a first-order autoregressive covariance (AR1) function (Cameletti et al., 2012). We set penalised complexity priors for the range and variance parameters of the spatial covariance function (Simpson et al., 2017). The Matérn (*α*=3/2) covariance function is commonly applied in spatial statistics to capture spatially smooth variation (as in Simmonds et al. (2020)). Priors were set to decrease the variation at relatively long distances (with a probability of 0.01 that the range parameter has a value corresponding to less than 200 km) and to constrain the magnitude of variance (with a probability of 0.1 that the standard deviation is greater than 0.5) following the recommendations by Fuglstad et al. (2018); Mäkinen et al. (2022). To avoid a scenario where spatio-temporal random effect explains all variation in the response variable and makes covariate effects redundant, we orthogonalised the spatio-temporal random effect to the covariates. Orthogonalisation has previously been successfully applied in Page et al. (2017); Hodges and Reich (2010).

The model intercepts and the effects of the environmental variables were assigned Gaussian distributed priors with a mean of zero and a variance of ten. To resolve the effect size of each variable and their interactions, we then examined the coefficient estimates of the model. Since we used scaled CTI values and covariates, the estimates of *β* reflect the change in scaled CTI with a change of one standard deviation of scaled covariate values. We fitted the full models for each taxa with INLA software in R (version 4.4.2; R Core Team (2021)) using the R-INLA package (version 24.12.11; Rue et al. (2009)). Outputs of the models are included in Supplementary Material (Tables S6-10).

#### Variance partitioning

The associations between CTI and environmental drivers capture the spatio-temporal dynamics. Since the study area covers a long climatic gradient from the hemiboreal bioclimatic zone to the southern edge of the arctic bioclimatic zone, the spatial gradients in CTI, temperature, and forest cover are considerable. Thus, these statistical associations may smudge the signal from the spatial and temporal drivers of CTI. To separate the importance of covariates and spatio-temporal random effect on the temporal change of CTI, we conducted a spatially-explicit variance partitioning in which we examined the impact of the realised change in environmental drivers on CTI separately for each monitoring site (Schulz et al., 2025). See spatial distributions of model outputs in Supplementary Material (Fig. S2).

#### Visualising habitat-dependent temperature sensitivity of CTI

To illustrate how habitat context modifies the relationship between temperature and changes in thermal community composition, we derived the temperature sensitivity of the Community Temperature Index (CTI) from the fitted models. In our model formulation,

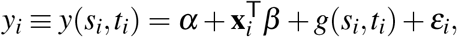

the vector of covariates **x**_*i*_ includes temperature (*T*), forest cover (*F*), landscape entropy (*E*), and the interactions between temperature and the two habitat context variables (*T × F* and *T × E*). The corresponding regression coefficients in *β* therefore quantify the effects of temperature (*β*_*T*_), forest cover (*β*_*F*_), entropy (*β*_*E*_), and the interaction terms (*β*_*TF*_ and *β*_*TE*_).

Expanding the relevant part of the linear predictor gives

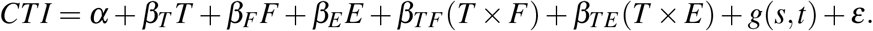

The expected change in CTI associated with temperature can then be obtained by taking the partial derivative of the linear predictor with respect to temperature,

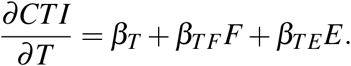

This expression represents the *temperature sensitivity of CTI*, that is, the expected change in CTI associated with a one-standard-deviation increase in temperature, conditional on habitat context.

To visualise how habitat structure modifies this temperature sensitivity, we evaluated the above expression across gradients of forest cover and landscape entropy. Habitat variables were varied across the observed range of standardised values (*−*2 to +2 standard deviations), while holding the other habitat variable constant at its mean. For each taxonomic group, predicted temperature sensitivity was therefore calculated as

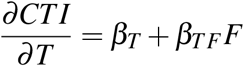

for the forest gradient, and

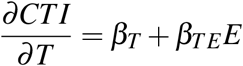

for the entropy gradient.

These predictions also provide a direct visualisation of the interaction terms in the model, illustrating how habitat context modifies the strength of the temperature–CTI relationship. Uncertainty bands were derived from posterior standard deviations of the corresponding regression coefficients.

## ACKNOWLEDGEMENTS

We thank all the volunteers and researchers who have collected and curated the data over several decades. We also thank the many people across Europe involved in the collation of species range maps to generate the Species Temperature Indexes. For moth range maps we thank Elisa Hanhirova and Ruby Fries for their time spent manually digitisation maps; for phytoplankton occurrence datasets we thank Wayne Trodd, Ute Mischke, Martin Søndergaard, Audron Pumputyt, Agnieszka Pasztaleniec, Christophe Laplace-Treyture, Gábor Borics, Aldo Marchetto, Otilia Mihail, Birger Skjelbred, Marko Järvinen, Stina Drakare, and Laurence Carvalho; for the plant range maps, we thank the ClimPlant team: Pieter Vangansbeke, František Máliš, Radim Hédl, Alistair Auffret, Jan Plue, Pieter De Frenne, and two additional sets of plant distribution maps provided by 1) Jesse Kalwij and 2) Erik Welk and Gunnar Seidler. We thank Pinja Kettunen for designing the conceptual Figure 1. The silhouette images were downloaded from www.phylopic.org and are by Andy Wilson (moth: *Arctia caja*, butterfly *Polygonia comma*), Sarah Frail (phytoplankton: *Thalassiosira pseudonana*) and others (plant: *Vaccinium erythrocarpum*). The bird (*Cyanistes caeruleus*) icon was manually vectorised by E.E.E. Finally, we thank Tanja Lindholm, Bess Hardwick, and Manuel Frias for their support in curating the species data, and recently retired Juha Pöyry of SYKE for curating the moth monitoring data. This work was funded by the Jane and Aatos Erkko Foundation (T.R., E.E.E., A.D., A.-L.L., M. Saastamoinen, I.C., E.K., J.V., and J.M.,). L.H.A. acknowledges funding from the Research Council of Finland (grant 361416/372215) and M.H.H. from the Research Council of Finland (grant 330739/360742). J.V. acknowledges funding from the European Union via ERC Consolidator Grant (BEFPREDICT, 101087409). A.S. was supported by a “Ramón y Cajal” fellowship (RYC2022-036239-I). The Butterfly Monitoring Scheme in Finnish Agricultural Landscapes (Diurna) was supported by the Finnish Ministry of the Environment. The data collection of understorey vegetation in Natural Resources Institute Finland (Luke) was conducted in the Biosoil project under the Forest Focus scheme (Regulation (EC) Nr. 2152/2003).

## DATA AVAILABILITY STATEMENT

All code to reproduce this analysis is available on Zenodo following this private peer review link xxx which will be made public on publication.

## Notes

### Competing Interest Statement

The authors have declared no competing interest.

