## Supplementary Material for "Habitat context alters the pace of climate-driven community warming across terrestrial and freshwater ecosystems"

[+]Equal contributions
[*]Corresponding author
[1]Research Center for Ecological Change, Department of Organismal and Evolutionary Biology, Faculty of Biological and Environmental Sciences, University of Helsinki, Helsinki, Finland
[2]Finnish Environment Institute (Syke), Helsinki, Finland
[3]Department of Biology, Faculty of Science, University of Turku, Finland
[4]Animal Demography and Ecology Unit, Institute for Mediterranean Studies (IMEDEA), CSIC-UIB, 07190, Esporles, Spain
[5]INRAE, EABX, 33612, Cestas, France
[6]Finnish Museum of Natural History, University of Helsinki, Finland
[7]Kainuu Centre for Economic Development, Transport and the Environment, Kajaani, Finland
[8]Natural Resources Institute Finland (Luke), Helsinki, Finland
[9]Department of Mathematics and Statistics, Faculty of Science, University of Helsinki, Helsinki, Finland
[10]Research Center for Ecological Change, Department of Environmental Science, Faculty of Biological and Environmental Sciences, University of Helsinki, Helsinki, Finland
[11]Department of Ecology, Swedish University of Agricultural Sciences, Uppsala, Sweden

### Supplementary Figures:

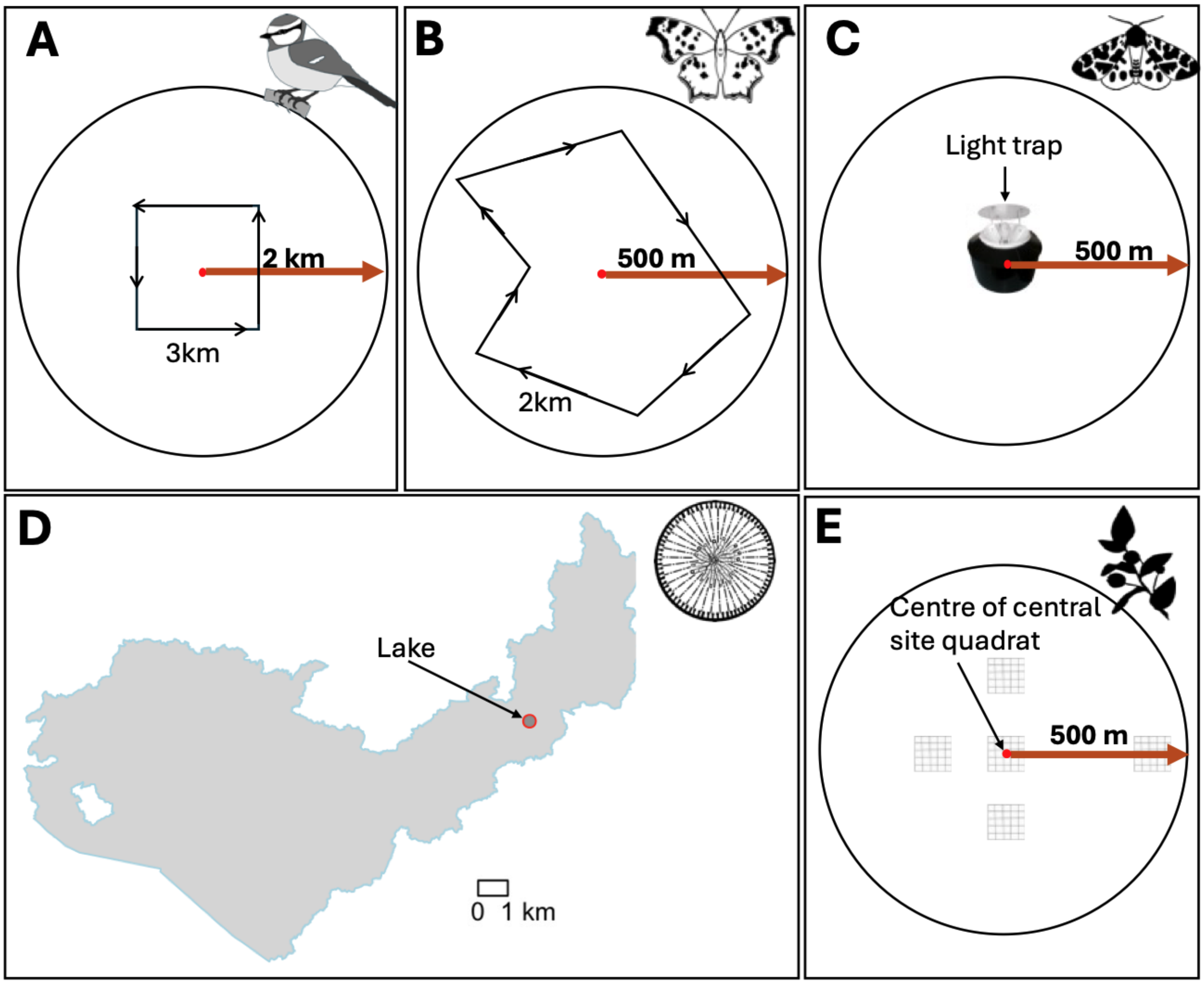


Figure S1: Illustrations of how we define taxon-specific buffers to extract habitat variables from. Buffers drawn around A) bird transects (2 km), B) butterfly transects (500 m), C) moth light traps (500 m), D) for phytoplankton, the catchment area surround the lake (catchment boundary), and E) the centre of the plant survey site (500 m). The buffer centre is defined a the centre of the transect (A,B), the light trap (C), the sampled lake within the catchment area (D) or the centre quadrat of the plant survey site (E)

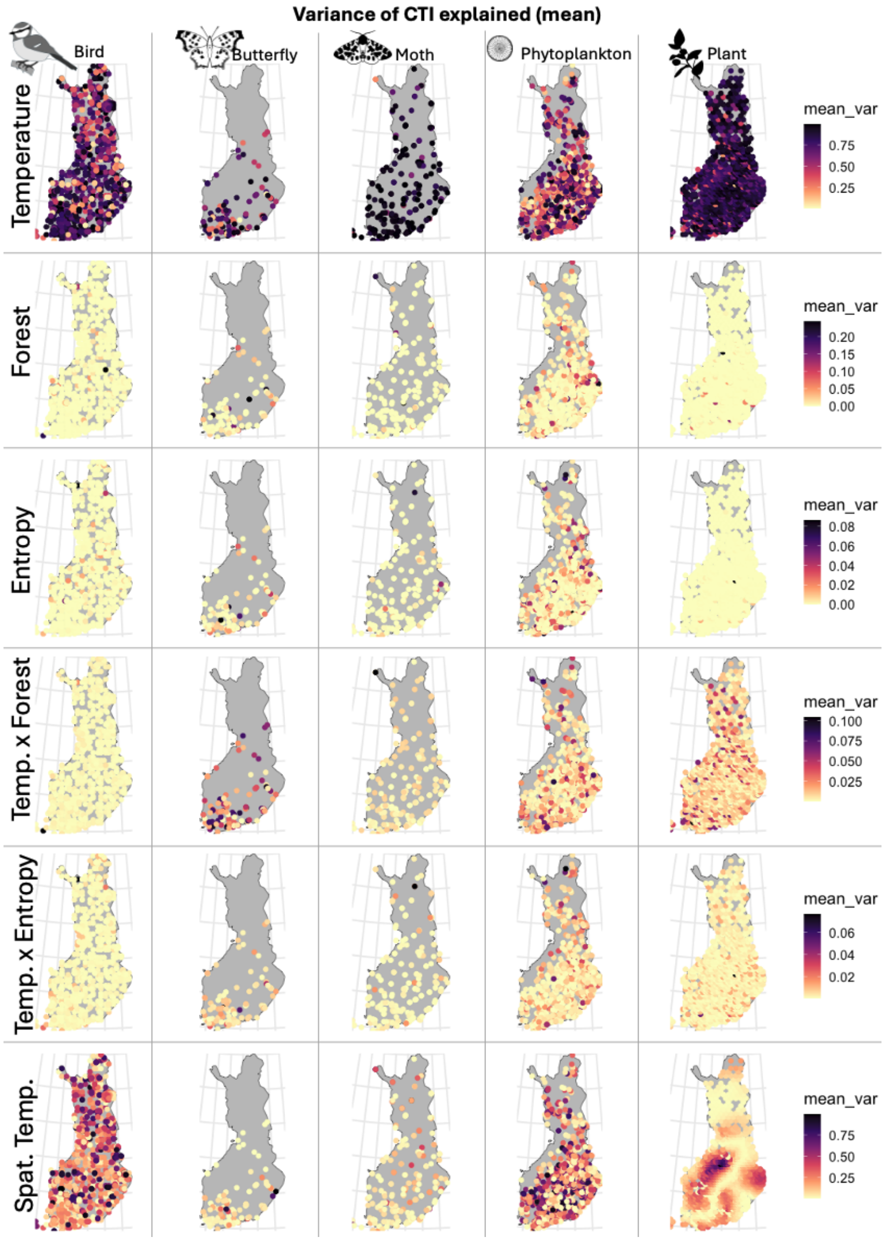


Figure S2: Spatial patterns in the variance of the Community Temperature Index (CTI) explained by different environmental predictors across Finland. Each column represents a taxonomic group (birds, butterflies, moths, phytoplankton, and plants), and each row shows the proportion of variance explained by a specific predictor in the variance partitioning analysis: temperature, forest cover, landscape entropy, the interaction between temperature and forest cover, the interaction between temperature and entropy, and spatial temperature structure. Points represent monitoring sites and are coloured by the mean proportion of variance explained by each predictor across years at that site (mean_var). Darker colours indicate higher explanatory power of the predictor for temporal variation in CTI at that location.

### Supplementary Tables

#### Environmental trends across monitoring sites

Table S1: Temporal trends in environmental covariates across moth monitoring sites. Site-level temporal trends ($\Delta$ per year) estimated from linear mixed models with site-specific random slopes for year. Values summarise the distribution of site-specific slopes across monitoring sites.

| Variable | Mean slope | SD | Median | 2.5% | 97.5% | N sites |
|---|---|---|---|---|---|---|
| Temperature ($^{\circ}$C yr$^{-1}$) | 0.0207 | 0.0261 | 0.0168 | -0.0349 | 0.0655 | 246 |
| Forest cover ($\Delta$ yr$^{-1}$) | 0.0012 | 0.0055 | 0.0001 | -0.0048 | 0.0145 | 246 |
| Habitat entropy ($\Delta$ yr$^{-1}$) | 0.0005 | 0.0067 | 0.0002 | -0.0131 | 0.0131 | 246 |

Table S2: Temporal trends in environmental covariates across bird monitoring sites. Site-level temporal trends ($\Delta$ per year) estimated from linear mixed models with site-specific random slopes for year. Values summarise the distribution of site-specific slopes across monitoring sites.

| Variable | Mean slope | SD | Median | 2.5% | 97.5% | N sites |
|---|---|---|---|---|---|---|
| Temperature ($^{\circ}$C yr$^{-1}$) | 0.0091 | 0.0207 | 0.0104 | -0.0272 | 0.0447 | 934 |
| Forest cover ($\Delta$ yr$^{-1}$) | 0.0009 | 0.0063 | 0.0001 | -0.0068 | 0.0180 | 934 |
| Habitat entropy ($\Delta$ yr$^{-1}$) | -0.0010 | 0.0081 | 0.0000 | -0.0184 | 0.0158 | 934 |

Table S3: Temporal trends in environmental covariates across plant monitoring sites. Site-level temporal trends ($\Delta$ per year) estimated from linear mixed models with site-specific random slopes for year. Values summarise the distribution of site-specific slopes across monitoring sites.

| Variable | Mean slope | SD | Median | 2.5% | 97.5% | N sites |
|---|---|---|---|---|---|---|
| Temperature ($^{\circ}$C yr$^{-1}$) | 0.1280 | 0.0385 | 0.1300 | 0.0668 | 0.1960 | 1538 |
| Forest cover ($\Delta$ yr$^{-1}$) | 0.0000 | 0.0017 | 0.0000 | -0.0030 | 0.0014 | 1538 |
| Habitat entropy ($\Delta$ yr$^{-1}$) | 0.0001 | 0.0025 | 0.0000 | -0.0032 | 0.0049 | 1538 |

Table S4: Temporal trends in environmental covariates across phytoplankton monitoring sites. Site-level temporal trends ($\Delta$ per year) estimated from linear mixed models with site-specific random slopes for year. Values summarise the distribution of site-specific slopes across monitoring sites.

| Variable | Mean slope | SD | Median | 2.5% | 97.5% | N sites |
|---|---|---|---|---|---|---|
| Temperature (°C $yr^{-1}$) | 0.0222 | 0.0218 | 0.0251 | -0.0266 | 0.0541 | 956 |
| Forest cover ($\Delta$ $yr^{-1}$) | 0.0011 | 0.0052 | 0.0004 | -0.0068 | 0.0130 | 956 |
| Habitat entropy ($\Delta$ $yr^{-1}$) | -0.0013 | 0.0062 | -0.0005 | -0.0146 | 0.0089 | 956 |

Table S5: Temporal trends in environmental covariates across butterfly monitoring sites. Site-level temporal trends ($\Delta$ per year) estimated from linear mixed models with site-specific random slopes for year. Values summarise the distribution of site-specific slopes across monitoring sites.

| Variable | Mean slope | SD | Median | 2.5% | 97.5% | N sites |
|---|---|---|---|---|---|---|
| Temperature (°C $yr^{-1}$) | 0.0171 | 0.0242 | 0.0144 | -0.0343 | 0.0633 | 101 |
| Forest cover ($\Delta$ $yr^{-1}$) | 0.0022 | 0.0101 | 0.0004 | -0.0123 | 0.0254 | 101 |
| Habitat entropy ($\Delta$ $yr^{-1}$) | -0.0020 | 0.0115 | -0.0002 | -0.0318 | 0.0159 | 101 |

### Posterior summaries of INLA models

Table S6: Fixed effects from INLA models for moth communities. Posterior summaries of fixed effects from Bayesian spatial models estimating drivers of temporal variation in the Community Temperature Index (CTI) across Finland.

| Predictor | Mean | SD | 2.5% CI | 97.5% CI |
|---|---|---|---|---|
| Intercept | -0.006 | 0.005 | -0.016 | 0.005 |
| Temperature | 0.807 | 0.011 | 0.786 | 0.828 |
| Forest cover | -0.186 | 0.011 | -0.207 | -0.165 |
| Landscape entropy | 0.019 | 0.011 | -0.003 | 0.040 |
| Temperature × Forest | 0.009 | 0.011 | -0.012 | 0.030 |
| Temperature × Entropy | 0.029 | 0.010 | 0.010 | 0.048 |

Table S7: Fixed effects from INLA models for butterfly communities. Posterior summaries of fixed effects from Bayesian spatial models estimating drivers of temporal variation in the Community Temperature Index (CTI) across Finland.

| Predictor | Mean | SD | 2.5% CI | 97.5% CI |
|---|---|---|---|---|
| Intercept | 0.018 | 0.013 | -0.008 | 0.043 |
| Temperature | 0.456 | 0.025 | 0.406 | 0.506 |
| Forest cover | -0.139 | 0.025 | -0.188 | -0.090 |
| Landscape entropy | -0.042 | 0.025 | -0.090 | 0.006 |
| Temperature × Forest | -0.177 | 0.025 | -0.225 | -0.128 |
| Temperature × Entropy | -0.038 | 0.028 | -0.094 | 0.017 |

Table S8: Fixed effects from INLA models for bird communities. Posterior summaries of fixed effects from Bayesian spatial models estimating drivers of temporal variation in the Community Temperature Index (CTI) across Finland.

| Predictor | Mean | SD | 2.5% CI | 97.5% CI |
|---|---|---|---|---|
| Intercept | 0.006 | 0.006 | -0.006 | 0.017 |
| Temperature | 0.709 | 0.012 | 0.685 | 0.732 |
| Forest cover | 0.069 | 0.010 | 0.050 | 0.088 |
| Landscape entropy | 0.032 | 0.010 | 0.013 | 0.051 |
| Temperature × Forest | -0.027 | 0.009 | -0.044 | -0.009 |
| Temperature × Entropy | -0.026 | 0.009 | -0.044 | -0.008 |

Table S9: Fixed effects from INLA models for plant communities. Posterior summaries of fixed effects from Bayesian spatial models estimating drivers of temporal variation in the Community Temperature Index (CTI) across Finland.

| Predictor | Mean | SD | 2.5% CI | 97.5% CI |
|---|---|---|---|---|
| Intercept | 0.001 | 0.008 | -0.014 | 0.016 |
| Temperature | 0.236 | 0.016 | 0.205 | 0.267 |
| Forest cover | -0.040 | 0.017 | -0.073 | -0.007 |
| Landscape entropy | 0.005 | 0.017 | -0.028 | 0.038 |
| Temperature × Forest | -0.068 | 0.016 | -0.099 | -0.037 |
| Temperature × Entropy | 0.035 | 0.016 | 0.003 | 0.067 |

Table S10: Fixed effects from INLA models for phytoplankton communities. Posterior summaries of fixed effects from Bayesian spatial models estimating drivers of temporal variation in the Community Temperature Index (CTI) across Finland.

| Predictor | Mean | SD | 2.5% CI | 97.5% CI |
|---|---|---|---|---|
| Intercept | 0.004 | 0.007 | -0.010 | 0.017 |
| Temperature | 0.243 | 0.014 | 0.216 | 0.269 |
| Forest cover | -0.260 | 0.014 | -0.287 | -0.233 |
| Landscape entropy | 0.207 | 0.014 | 0.180 | 0.233 |
| Temperature × Forest | -0.086 | 0.014 | -0.113 | -0.058 |
| Temperature × Entropy | -0.002 | 0.013 | -0.029 | 0.024 |
